# Compression of redundant visual information improves feature discrimination in human vision

**DOI:** 10.64898/2026.09.07.749904

**Authors:** Li L-Miao, Doğukan Nami Öztaş, Nihan Alp, Bilge Sayim

## Abstract

The visual environment contains more information than can be fully processed. To cope with this information, the visual system selects, integrates, and compresses sensory inputs. These processes are particularly evident in peripheral vision, where nearby items strongly limit access to individual elements, as in crowding and redundancy masking. In redundancy masking, repeated items are compressed into fewer items than are physically present, for example, three spatially separated items presented in the periphery are often perceived as only two items. Whether these interactions among items merely limit access to visual information or serve a functional role in perception remains unknown. Here, we investigated whether redundancy masking, despite reducing the number of perceived items, improves feature discrimination. In two experiments, participants viewed arrays of identical bars in peripheral vision, reported the number of bars they perceived, and adjusted a foveal probe to match perceived width (Experiment 1) or width and spacing (Experiment 2). Strong redundancy masking occurred in both experiments. When participants perceived fewer bars than were presented, perceived width was closer to the presented bar width than when all bars were perceived, where width was systematically underestimated. In Experiment 2, the perceived spacing between the (fewer) perceived bars increased. We modeled feature discrimination under redundancy masking and found that the observed improvement could not be explained by averaging features of the lost and perceived bars. These results suggest that the visual system efficiently compresses redundant information, reducing the number of perceived items while improving feature discrimination.

## INTRODUCTION

The visual environment contains far more information than the human visual system can fully process. To cope with this complexity, the visual system selectively reduces information through processes such as selection, integration, and compression of sensory inputs. One way to reduce the amount of information that needs to be processed is to identify redundancies in the visual input and represent them in a more compact format. Textures provide a canonical example because each element shares much of its information with its neighbours, and the visual system can represent the statistical regularities of textures without encoding every element individually (Julesz, 1981; Simoncelli & Portilla 1998; Portilla & Simoncelli 2000; Rosenholtz, 2014). This is a form of compression in which redundant structure is represented in a more economical form (e.g., Barlow, 1961). Importantly, such compression is particularly pronounced in peripheral vision, where the visual system represents information with lower spatial precision (Anderson et al., 1991; Strasburger et al., 2011), and relies more heavily on statistical regularities in the visual environment (Chetverikov et al., 2016; Fiser & Aslin, 2001; Turk-Browne et al., 2009). Compression is generally thought to increase efficiency (Barlow, 1961; Simoncelli & Olshausen, 2001) and reduce processing demands (Attneave, 1954; Laughlin, 2001; Lennie, 2003; Rosenholtz, 2016) at the expense of visual detail (Balas et al., 2009; Bates & Jacobs, 2020; Rosenholtz et al., 2012; Zhaoping, 2006). Whether this reduction in visual detail is solely detrimental, or can also benefit perceptual performance, remains an open question.

For texture perception, the loss of information about individual elements constituting the texture has little perceptual cost because the contribution of an individual element is largely predictable by its neighbors. This compression becomes problematic, however, when individual items must be detected or discriminated. One form of interaction between neighboring items is visual crowding. When a peripheral target is presented among nearby flankers, the flankers interfere with target perception (Bouma, 1970; Kooi et al., 1994; Pelli et al., 2004; Whitney & Levi, 2011; Herzog et al., 2015; Sayim & Rummens, 2026). Crowding is a major bottleneck of object recognition in peripheral vision (Levi, 2008; Pelli & Tillman, 2008; Whitney & Levi, 2011).

Crowding impairs target discrimination and distorts target appearance, but it does not interfere with target detection (Pelli et al., 2004; Pelli & Tillman, 2008; Whitney & Levi, 2011; Levi et al., 2002). A special case arises when the target and its flankers are identical. In this case, standard identification performance is at ceiling: for example, a target letter T flanked by two identical Ts in the visual periphery is reported correctly on almost every trial (Sayim & Taylor, 2019). However, when participants freely verbally report or draw what they perceive, they typically report only two Ts instead of three (Sayim & Taylor, 2019). Observers fail to report an entire item, a phenomenon known as redundancy masking (Sayim & Taylor, 2019; Yildirim et al., 2020, 2021, 2022). The same physical displays thus produce different outcomes depending on the task. Redundancy masking is a form of compression of visual information: multiple identical items in the input are represented by fewer items in the percept. This compression can be substantial: when three items are reported as two, one-third of the items go unreported.

Compression is usually assumed to trade visual detail for efficiency (Balas et al., 2009; Bates & Jacobs, 2020; Rosenholtz et al., 2012; Zhaoping, 2006), and reduce processing demands (Yildirim & Sayim, 2022). When an item is identical to its neighbors, however, compression can remove that item without discarding feature information that is unique to it, because its features do not differ from the surrounding items. This raises a broader question about what happens to the information that remains perceptually available. Redundancy masking could simply remove items while leaving the representation of the other items unchanged, or it could alter the representation of the remaining items. Previous EEG frequency tagging results suggest that redundancy masking does not eliminate the neural representation of the masked item: neural responses to all presented items persisted even when one item was redundancy-masked, and the loss of an item was accompanied by stronger neural integration between neighbouring items (Oztas et al., 2025). These results suggest that information from the lost item may still be represented in the neural responses to its neighbors, despite the item not being consciously perceived. If the remaining items carry the information of the item that was lost, their features should be represented differently than when every item is perceived. Here, we asked whether compression of redundant information can benefit perception: Are the features of the perceived items judged more accurately under redundancy masking than when the full set of items is perceived?

To address this question, we conducted two experiments to measure how redundancy masking changes perceived bar widths. In both experiments, participants viewed arrays of 3 to 5 identical bars in peripheral vision. They reported the number of bars they perceived and then adjusted a probe to match the perceived bar width. In Experiment 1, the probe was a single bar, isolating width perception. In Experiment 2, the probe contained the reported number of bars, and participants adjusted both the width and spacing between the bars, so that each trial yielded the perceived spatial configuration of the array. This also allowed us to assess the perceived extent of the array and the proportion of the extent covered by bars (density). Crucially, the same stimuli could yield redundancy-masking (RM) when participants reported fewer items than were presented (RM trials) and no redundancy masking when they reported all items that were presented, or more (no-RM trials). The perceived width of the remaining bars can therefore reveal how the width of the unreported bar is reflected in the perception of the remaining bars. One possibility is that the width of the unreported bar is pooled with the widths of the perceived bars. Pooling is one account of contextual interactions in peripheral vision, in which feature signals from an item and its neighbours are combined across a spatial region (Parkes et al., 2001; Greenwood et al., 2009; van den Berg et al., 2010). Our displays provide a direct test of pooling models. If the width of the redundancy-masked bar were completely pooled with the perceived bars, the total width of the array would be preserved, and the perceived bars would be proportionally wider than the presented bars with the magnitude of the increase determined by the ratio of presented to perceived bars. If the width of the redundancy-masked bar were not pooled, the width estimate should be unchanged by redundancy-masking. Values between these predictions would be consistent with partial pooling.

Both experiments showed strong redundancy masking, with participants frequently reporting fewer bars than were presented. When a bar was redundancy-masked, the widths of the perceived bars were judged more accurately than when all bars were reported. Computational modeling separated the general underestimation of width in peripheral vision from the contribution of the redundancy-masked bar and showed that most of its width was lost with the item rather than pooled into the reported bars. In Experiment 2, redundancy masking was also related to systematic changes in the spatial configuration of the array: the overall extent was underestimated and the spacing between perceived bars was overestimated, while density was much less affected. Together, these results show that the visual system does not simply lose redundant information but compresses it, reducing the number of perceived items while improving the feature discrimination of those that remain.

## Experiment 1

### METHOD

#### Participants

Twenty-one participants (2 males and 19 females, mean age: 22.3 years, range: 18 to 31) from Sabanci University participated in exchange for course credits. All participants were naive to the task and had normal or corrected-to-normal visual acuity. Prior to the experiment, participants were informed about the procedure and signed a consent form. The experimental protocol was approved by the Ethics Committee of Sabanci University. We ran a simulation-based power analysis with 1000 simulations per condition prior to data collection (Kumle et al., 2021). The power for our primary effect of interest (the main effect of RM type (RM trials, no-RM trials) on width perception, see Design and Procedure) reached 0.9 with a sample size of 5 participants.

#### Apparatus and Stimuli

The experiment was programmed in PsychoPy3 Builder (v2023.1.1; Peirce et al., 2019) and ran on a desktop PC with a 120 Hz monitor. The experiment was conducted in a noise- and distraction-free cubicle containing a desk, a non-wheeled chair, and a desktop computer. Viewing distance was maintained at 57 cm. Stimuli were arrays of 3 to 5 identical black bars presented on a gray background. Bar width was 0.25°, 0.4°, or 0.55°, and bar height was fixed at 1.9°. The center-to-center spacing was 0.7°, 0.9°, or 1.1°, with spacing adjusted across set sizes (3-bar arrays: 1.4°, 1.48°, 1.8°, 2.2°; 4-bar arrays: 0.47°, 0.60°, 0.73°, 0.78°; 5-bar arrays: 0.53°, 0.68°, 0.83°, 0.86°) to keep the overall array extent constant across trials. This prevented participants from inferring the number of bars from the total array extent. For example, three bars of width 0.25° separated by 0.7° yielded a total array extent of 1.65°. This length was matched for four- and five-bar configurations by reducing the center-to-center spacing between adjacent bars to 0.47° and 0.35°, respectively. Set sizes, widths, and spacings were selected based on prior work showing that these values are optimal for redundancy masking (Yildirim et al., 2020).

#### Design and Procedure

The experiment used a within-subjects design crossing set size (3, 4, 5), bar width (0.25°, 0.4°, 0.55°), and spacing (0.7°, 0.9°, 1.1°). For each combination of set size, width, and spacing, we included the same number of trials with matched spacing, with spacing adjusted to keep the overall array extent constant across set sizes. This yielded 54 trials (3 set sizes × 3 widths × 3 spacings × 2) per block. The experiment consisted of 10 blocks.

Participants received written and verbal instructions and completed 10 practice trials before the main experiment. Participants were not informed about the possible number of bars presented in the experiment. On each trial, a black fixation dot (diameter: 0.11°) was presented at the center of the screen for a randomly selected duration of 1, 1.25, 1.5, 1.75, or 2 seconds. An array of 3 to 5 identical black bars was then presented for 150 ms, either to the left or to the right of the fixation at 10° eccentricity. Participants first reported the number of bars they perceived by pressing the corresponding key on the numerical keypad. Single-digit responses from 0 to 9 were allowed. After the number report, a single probe bar appeared at the center of the screen. Participants adjusted the probe width to match the perceived width of the bars presented. For half of the participants, width was adjusted with the left and right arrow keys; for the other half, it was adjusted with the up and down arrow keys. Each key press increased or decreased the width by 0.05°. The initial probe width was randomly selected from a uniform distribution ranging from 0.2° to 0.6° in increments of 0.05°. Participants pressed the “Enter” key to confirm their adjustment and proceed to the next trial. A schematic depiction of the trial sequence is shown in Figure 1A.

**Figure 1.**
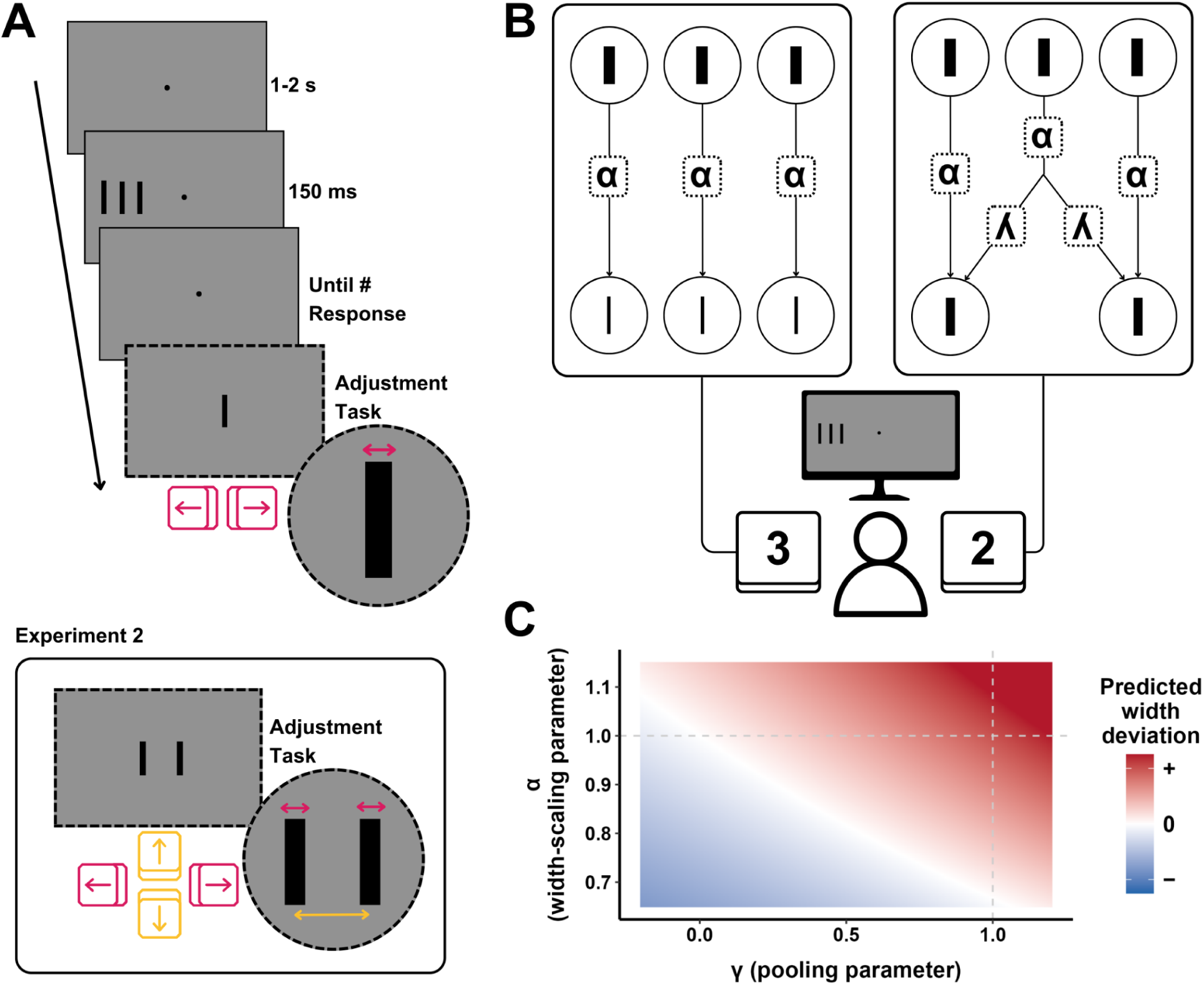
Experimental procedure and computational models. (A) Trial sequence (see Methods for details). An array of 3–5 identical bars appeared for 150 ms, and participants reported the perceived number of bars, then matched a foveal probe to the perceived bar width (magenta). In Experiment 2 (bottom), the probe contained the reported number of bars, and participants also adjusted inter-bar spacing (yellow). (B) Model schematic. On no-RM trials (left), each bar is scaled by the width-scaling parameter α, and the perceived number equals the presented number. On RM trials (right), α is followed by a pooling stage (parameter γ) in which features are pooled across adjacent bars, and fewer bars are perceived. (C) Width deviation predicted by M5 (Linear width-scaling + Partial Pooling) as a function of γ (x-axis) and α(W) = α₀ + α₁W (y-axis): blue = underestimation, white = veridical, red = overestimation. Dashed lines mark α = 1 and γ = 1; the example is the 3→2 RM trial at W = 0.4°.

After the main experiment, participants completed a single-bar control task, providing a baseline for peripheral width perception in the absence of neighboring bars. Here, a single bar was presented at 10° eccentricity, and participants matched a foveal probe to its perceived width (30 trials).

#### Analysis

All analyses were conducted using R (R Core Team, 2024). Number deviation was computed as the presented subtracted from the reported number of bars, and width deviation as the presented subtracted from the reported width. Trials were excluded if response times exceeded 10 seconds or number deviation fell outside +/- 4 bars in the enumeration tasks (Yildirim & Sayim, 2022), or if response time exceeded 15 seconds in the adjustment task. In total, 0.06% of trials were removed. For comparing RM and no-RM trials, analyses were restricted to trials with number deviations of −1 (RM) and 0 (no-RM). Trials with under-reporting of multiple bars and over-reporting were excluded because they reflect qualitatively different responses. This exclusion removed 12% of trials (see Table S1 for the full distribution of number deviations).

Linear mixed-effects models were fitted using the lme4 package in R (Bates et al., 2015) with width deviation as the dependent variable and RM type, set size (3, 4, 5), bar width (0.25°, 0.4°, 0.55°), and all interactions as fixed effects, with by-participant random intercepts and random slopes for RM type. Post-hoc RM vs. no-RM contrasts were computed using the emmeans package (Lenth, 2021) using FDR correction (Benjamini & Hochberg, 1995). Standardized effect sizes were calculated by dividing contrast estimates by the model’s residual standard deviation.

To assess width perception, we first computed relative width deviation for each trial by dividing the absolute width deviation by the presented width. One-sample t-tests compared these relative deviations against zero (no deviation), separately for RM, no-RM, and single-bar trials. Paired comparisons further compared RM against no-RM trials, and each against the single-bar condition.. P-values were adjusted using the FDR correction, and effect sizes are reported as Cohen’s d, computed as the mean difference divided by the standard deviation of the differences.

For each participant, we quantified the precision of width perception as the standard deviation (SD) of width deviations, computed separately for each RM type and each presented width. Precision was compared across conditions using three paired t-tests (RM vs. no-RM, RM vs. single-bar, no-RM vs. single-bar; FDR-corrected; see Supplementary Section S1 for details) To examine whether the perceived width depended on the presence or absence of RM, we fitted a computational model to each participant’s trial-level width reports. The model tested to what extent the width of the redundancy-masked bar was lost or pooled into the perceived/remaining bars, predicting perceived width (W′), from presented width (W), using two parameters. The first, width-scaling (α), captures underestimation of target width in peripheral vision (Newsome, 1972; Baldwin et al., 2016). The second, pooling (γ), captures the extent to which a redundancy-masked bar’s width is pooled into the perceived width of the reported bars. Pooling is determined by the ratio of presented to perceived number of bars (N/N′), which equals 1 for no-RM trials and exceeds 1 when redundancy masking occurs (Figure 1B). γ ranged from no pooling (γ = 0) to full pooling (γ = 1).

Width-scaling linearly varies with bar width:

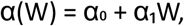

so width estimation can differ across the presented widths. The two effects combine as

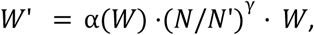

where the presented width W is scaled by α(W) (width underestimation), and multiplied by the pool ratio raised to γ (pooling) to give the predicted width W′ (Figure 1C). Hence, the model separates two sources of perceived width: general peripheral width underestimation and the potential pooling of information from a redundancy-masked item into the perceived items. We refer to this model as M5 (Linear Width-Scaling + Partial Pooling). It provided the best fit among a set of models that progressively added width-scaling and pooling to a null baseline, fit per participant by minimizing squared error and compared by AIC and likelihood ratio tests. Full specifications of all models, derivations, and fitting details are in the Supplementary Material (Section S2).

### Results

#### Number Deviation

Participants reported fewer bars than presented, indicating strong redundancy masking (Figure 2A). The proportion of trials in which the number of bars was underestimated increased with set size: 26.9% for set size 3, 37.6% for set size 4, and 40.9% for set size 5. Mean number deviations were −0.18 (SD = 0.33) for set size 3, −0.33 (SD = 0.39) for set size 4, and −0.34 (SD = 0.56) for set size 5.

**Figure 2.**
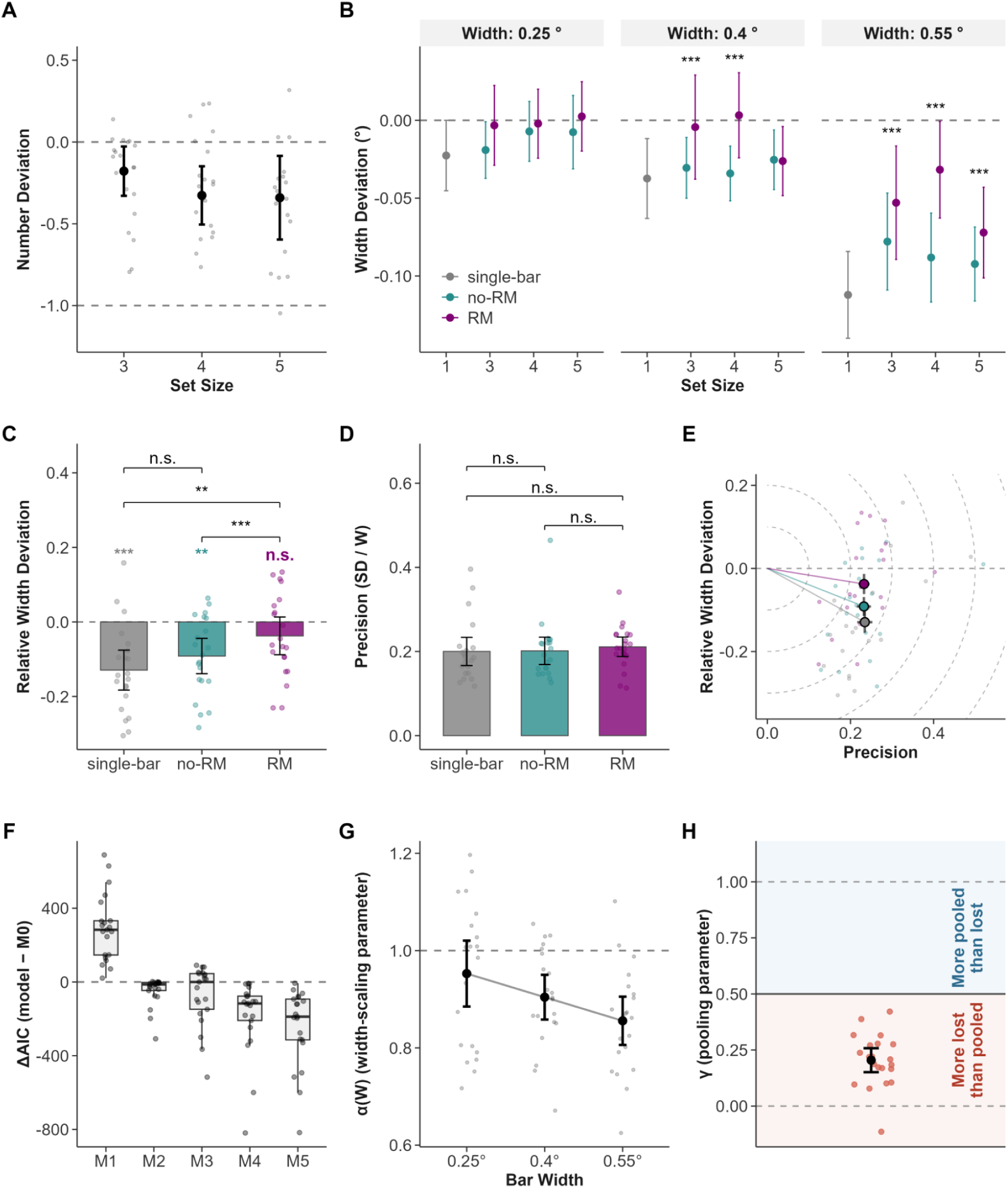
Experiment 1 results (n = 21 participants). Dots are participant means, error bars 95% CIs; asterisks mark significant RM vs no-RM contrasts (*p < .05; **p < .01; ***p < .001). (A) Number deviation by set size. (B) Width deviation by set size and presented width. (C) Relative width deviation for single-bar, no-RM, and RM; asterisks here mark deviation from the zero dashed line. (D) Precision by condition; brackets show FDR-corrected pairwise comparisons. (E) Joint deviation–precision space; large circles are group means, colored segments from the origin show RMSE. (F) Model comparison (ΔAIC relative to M0). (G) Width-scaling α(W) by bar width (M5); dashed line at α = 1. (H) Pooling parameter γ (M5); dashed lines at 0 and 1, solid line at 0.5; red/blue split participants below vs above 0.5.

#### Width Deviation

To test whether the loss of conscious access to an item impaired or improved feature perception, we compared perceived bar width across RM and no-RM trials. Overall, perceived bar width was greater in RM trials than in no-RM trials, where width was systematically underestimated (Figure 2B; trial percentages per condition in Table S2). Significant RM effects were observed for medium-width bars (0.40°) at set sizes 3 (β = −0.031, z = −4.82, d = 0.37, 95% CI [−0.043, −0.018]) and 4 (β = −0.033, z = −5.40, d = 0.39, 95% CI [−0.045, −0.021]), and for large bars (0.55°) at all set sizes (β = −0.024, z = −4.01, d = 0.29, 95% CI [−0.036, −0.012], all p < .001). No significant RM effects were observed for narrow bars (0.25°) at any set size (all p ≥ .05), or for 0.40° bars at set size 5 (p = .29).

#### Width Deviation Compared to Zero

Relative width deviation was closer to zero on RM than on no-RM trials (t(20) = 6.67, p < .001, d = 1.46, 95% CI [0.037, 0.071]), with 20 of 21 participants showing this pattern. Significant width underestimation was found for no-RM trials (t(20) = −4.03, p = .001, d = −0.88), and in the single-bar condition (t(20) = −5.03, p < .001, d = −1.10). By contrast, RM trials did not significantly differ from zero (t(20) = −1.54, p = .138, d = −0.34), suggesting that width perception in RM trials was close to veridical (Figure 2C).

#### Width Deviation Compared to Single-Bar

RM trials also showed significantly wider perceived widths than single-bar trials (t(20) = 3.99, p = .001, d = 0.87). No-RM trials did not significantly differ from the single-bar (t(20) = 1.55, p = .138, d = 0.34; Figure 2C). This result suggests that RM improved width discrimination, even compared to single bars.

#### Precision

Precision was similar across conditions (Figure 2D, Supplementary Section S1): single-bar M = 0.200 (SD = 0.074), no-RM M = 0.202 (SD = 0.072), and RM M = 0.211 (SD = 0.051). Paired comparisons (FDR-corrected across three tests) revealed no differences between RM and no-RM (t(20) = 1.01, p = .680, d = 0.22), RM and single-bar (t(20) = 0.76, p = .680, d = 0.17), or no-RM and single-bar (t(20) = 0.08, p = .937, d = 0.02). Additionally, Figure 2E shows width deviation plotted against precision. The root-mean-square error (RMSE), the total error combining deviation and precision, was numerically lowest on RM trials (RMSE = 0.26 for RM, 0.27 for no-RM, 0.29 for single-bar).

#### Modelling Results

To determine whether the lost item’s features were pooled into the perceived items, we compared computational models of perceived width. The Linear Width-scaling + Partial Pooling model (M5) provided the best fit (AIC; 18 of 21 participants), and likelihood ratio tests confirmed that each added parameter improved fit (Tables S3 and S4; Figure 2F). In M5, width-scaling was below 1 and decreased with bar width (α₀ = 1.03, SD = 0.27; α₁ = −0.32, SD = 0.55), reproducing width underestimation (Figure 2G). Pooling, however, was weak: the pooling parameter fell below 0.5 for every participant (M = 0.20, SD = 0.12; Figure 2H), indicating that the width of the redundancy-masked bar was not pooled into the perceived bars (Table S5).

## Experiment 2

### METHOD

#### Participants

Twenty participants (2 males and 18 females, mean age: 21.65 years, ranging from 19 to 24) from Sabanci University participated for course credits. Recruitment, consent, ethics approval, and the simulation-based power analysis followed the same procedures as Experiment 1.

#### Apparatus and Stimuli

Apparatus and stimuli used in Experiment 2 were identical to those in Experiment 1.

#### Design and Procedure

The design and procedure were identical to Experiment 1, except for the adjustment task. After reporting the number of bars, participants saw a probe array containing that same number of bars and adjusted both its width and its inter-bar spacing to match the perceived stimulus (Figure 1A). All bars in the probe changed together, so it was always uniform in width and evenly spaced. Width and spacing were each controlled by one pair of arrow keys, counterbalanced across participants, with each key press changing width or spacing by 0.05°. The probe’s initial width was drawn from a uniform distribution over 0.2°–0.6° (0.05° steps), and its initial spacing from a uniform distribution over 0.3°–1.2° (0.1° steps). The experiment comprised six blocks of 54 trials (324 per participant).

As in Experiment 1, participants then completed the single-bar control task, here with 18 trials (three widths). A coding error left two participants without single-bar data, so all single-bar comparisons in Experiment 2 are based on the remaining 18 participants (Supplementary Section S3). Participants also completed a two-bar discrimination control confirming that the bars were spatially resolvable at the eccentricities and spacings used in the main experiment (see Supplementary Section S3).

#### Analysis

Data analysis followed the same procedures used in Experiment 1. In addition to number and width deviation, we computed spacing deviation as the reported minus the presented spacing and, for each trial, relative spacing deviation as the spacing deviation divided by the presented spacing. Because length-matched trials produced a wide range of center-to-center spacings that depended on the corresponding width and set size (see Experiment 1, Apparatus and Stimuli), spacings were grouped into three categories for analysis: smaller (≤ 0.8°), middle ( > 0.8° and ≤ 1.0°), and larger ( > 1.0°). The exclusion criteria were identical to those in Experiment 1: 0.54% of trials were removed in total. For the RM vs. no-RM contrasts, the dataset was further restricted to trials with a number deviation of −1 or 0, corresponding to RM and no-RM trials, respectively. This restriction excluded an additional 8.02% of the remaining trials (Table S1).

Width and spacing deviations were analyzed separately using linear mixed-effects models. Each model included RM type, spacing category (smaller, middle, larger), set size (3, 4, 5), and bar width (0.25°, 0.4°, 0.55°) as fixed effects with all interactions, by-participant random intercepts, and random slopes for RM type. For width deviation, post-hoc RM vs. no-RM contrasts were computed with emmeans, conditioned on bar width and set size; for spacing deviation, contrasts were conditioned on spacing category and set size. Relative spacing deviation was computed per trial as the spacing deviation divided by the presented spacing. To test whether the RM effect on perceived width was associated with the RM effect on perceived spacing across participants, we computed, for each participant, Δwidth (mean relative width deviation on RM trials minus that on no-RM trials and Δspacing (the corresponding difference for relative spacing deviation), and correlated the two (Pearson).

We additionally computed two array-level measures. The perceived array extent (E′) was the edge-to-edge horizontal extent of the reproduced array,

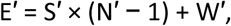

where S′, N′, and W′ are the perceived spacing, perceived number of bars, and perceived width, respectively. The array extent deviation was E′ − E, the perceived minus the presented array extent (E = S × (N − 1) + W, where S, N, and W are the presented spacing, number, and width), with negative values indicating a radially underestimated array. The perceived density (D′) was the summed bar width divided by the array extent,

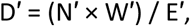

that is, the proportion of the array covered by bars. The density deviation was D′ − D, the perceived minus the presented density (D = (N × W) / E), with positive values indicating a denser percept. Array extent and density deviations were each analyzed with the same full-factorial linear mixed-effects model used for the width and spacing deviations, including RM type, spacing category (smaller, middle, larger), set size (3, 4, 5), and bar width (0.25°, 0.4°, 0.55°) with all interactions. The density model included participant-by-participant random intercepts and random slopes for RM type; for array extent, the RM type random slope produced a singular fit and was not supported by a likelihood-ratio test, so a random-intercept-only structure was used. Post-hoc RM vs. no-RM contrasts were computed with emmeans conditioned on set size, FDR-corrected, with Cohen’s d computed as the contrast estimate divided by the model’s residual standard deviation.

### RESULTS

#### Number Deviation

Participants reported fewer bars than presented, indicating strong redundancy masking. The proportion of trials in which the number of bars was underestimated increased with set size: 20.4% of trials for set size 3, 30.8% for set size 4, and 54.1% for set size 5. Mean number deviation was −0.15 (SD = 0.26) for set size 3, −0.25 (SD = 0.30) for set size 4, and −0.50 (SD = 0.46) for set size 5 (Figure 3A).

**Figure 3.**
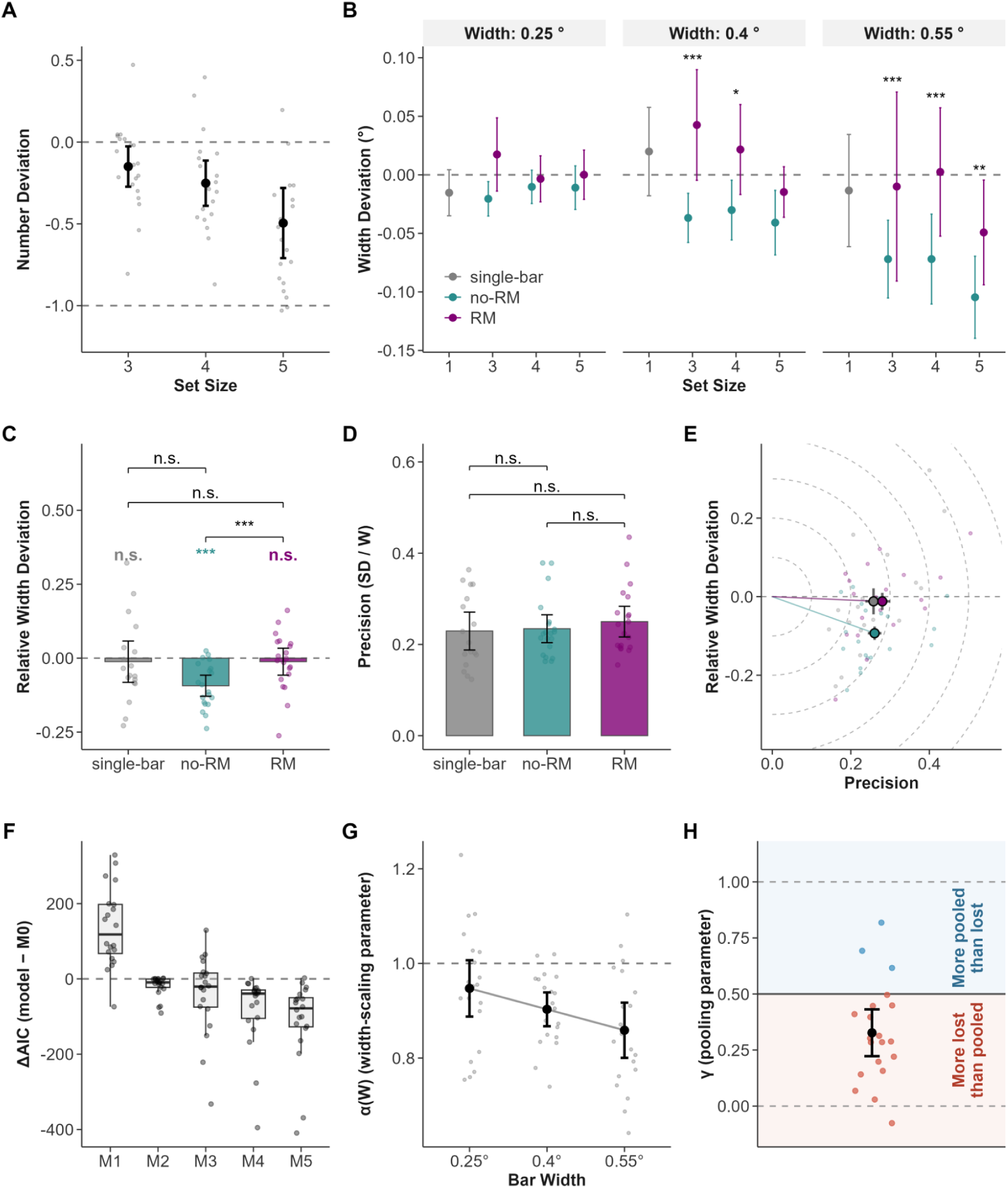
Experiment 2 results (n = 20 participants). Panels and conventions as in Figure 2.

#### Width Deviation

Significant RM effects were observed for: (a) 0.40° width at set sizes 3 (β = −0.067, z = −5.15, p < .001, d = 0.66, 95% CI [−0.093, −0.042]) and 4 (β = −0.033, z = −2.47, p = .024, d = 0.32, 95% CI [−0.059, −0.007]); and (b) 0.55° width at all set sizes (set size 3: β = −0.094, z = −7.73, p < .001, d = 0.91, 95% CI [−0.117, −0.070]; set size 4: β = −0.049, z = −4.31, p < .001, d = 0.48, 95% CI [−0.072, −0.027]; set size 5: β = −0.034, z = −3.15, p = .004, d = 0.33, 95% CI [−0.055, −0.013]). No significant RM effects were found for 0.25° width bars at any set size (all p ≥ .27). Overall, perceived width was greater in RM trials than in no-RM trials, indicating that bars appeared wider when RM occurred (Figure 3B; trial percentages per condition in Table S2).

#### Width Deviation Compared to Zero

Relative width deviation was closer to zero on RM than on no-RM trials (t(19) = 6.30, p < .001, d = 1.41, 95% CI [0.054, 0.108]), with 19 of 20 participants showing this pattern. No-RM trials showed significant underestimation (t(19) = −5.47, p < .001, d = −1.22). RM trials did not significantly differ from zero (t(19) = −0.55, p = .868, d = −0.12), indicating that width perception in RM trials was close to veridical (Figure 3C). The single-bar also did not significantly differ from zero (t(17) = −0.36, p = .868, d = −0.08).

#### Width Deviation Compared to Single-Bar

Neither RM (t(17) = 0.14, p = .889, d = 0.03) nor no-RM (t(17) = −2.40, p = .057, d = −0.57) trials significantly differed from the single-bar (Figure 3C).

#### Precision

Precision was similar across conditions: single-bar M = 0.229 (SD = 0.081), no-RM M = 0.234 (SD = 0.065), and RM M = 0.251 (SD = 0.071). FDR-corrected paired comparisons revealed no significant differences between RM and no-RM (t(16) = 2.02, p = .180, d = 0.49), RM and single-bar (t(16) = 1.14, p = .407, d = 0.28), or no-RM and single-bar (t(16) = 0.31, p = .761, d = 0.08; Figure 3D; Supplementary Section S1). Figure 3E shows width deviation plotted against precision. RM trials and the single-bar were both close to zero deviation, whereas no-RM trials shifted away from zero, and RMSE is similar across all three conditions (RMSE = 0.30 for RM, 0.29 for no-RM, 0.29 for single-bar).

#### Modelling Results

As in Experiment 1, the Linear Width-scaling + Partial Pooling model (M5) provided the best fit (AIC; 15 of 20 participants), and likelihood ratio tests again confirmed that each added parameter improved fit (Tables S3 and S4; Figure 3F). Width-scaling was below 1 and decreased with bar width (α₀ = 1.02, SD = 0.28; α₁ = −0.30, SD = 0.67; Figure 3G), and pooling was again weak: the pooling parameter fell below 0.5 for most participants (M = 0.32, SD = 0.22; 17 of 20; Figure 3H), so most of a lost bar’s width was not pooled into the remaining bars (Table S5).

#### Spacing Deviation

Perceived spacing was larger in RM than in no-RM trials in all nine combinations of set size and spacing category (all p < .01, FDR-corrected; Figure 4A). At set size 3, the RM vs. no-RM contrast was β = −0.123 (z = −3.05, d = 0.45, 95% CI [−0.203, −0.044]) for smaller, β = −0.251 (z = −6.26, d = 0.91, 95% CI [−0.330, −0.173]) for middle, and β = −0.641 (z = −22.45, d = 2.33, 95% CI [−0.697, −0.585]) for larger spacing. At set size 4, the contrast was β = −0.138 (z = −4.89, d = 0.50, 95% CI [−0.193, −0.083]) for smaller, β = −0.134 (z = −3.83, d = 0.49, 95% CI [−0.203, −0.066]) for middle, and β = −0.135 (z = −3.27, d = 0.49, 95% CI [−0.217, −0.054]) for larger spacing. At set size 5, the contrast was β = −0.098 (z = −3.45, d = 0.36, 95% CI [−0.154, −0.042]) for smaller, β = −0.135 (z = −4.29, d = 0.49, 95% CI [−0.197, −0.074]) for middle, and β = −0.196 (z = −5.31, d = 0.72, 95% CI [−0.269, −0.124]) for larger spacing. Thus, redundancy masking consistently expanded the perceived spacing between bars (trial percentages per condition in Table S2).

**Figure 4.**
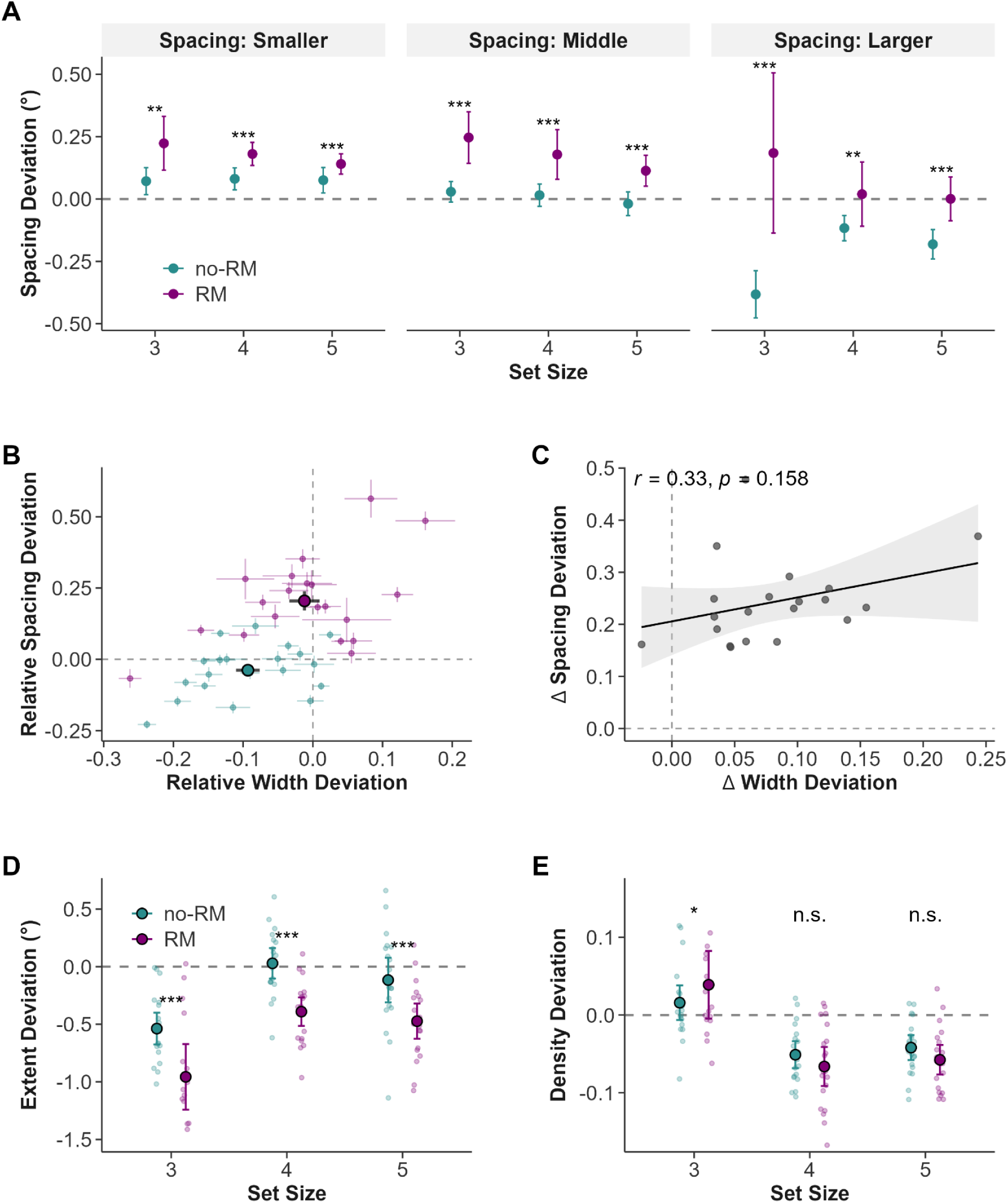
Spacing, extent, and density in Experiment 2. Conventions as in Figure 2. (A) Spacing deviation by set size and spacing category. (B) Joint width- × spacing-deviation space; large circles are group means. (C) Across-participant relationship between RM effects: Δwidth (RM − no-RM) vs Δspacing (RM – no-RM), one dot per participant. (D) Array-extent deviation by set size; negative values indicate underestimated extent (E) Density deviation by set size.

#### Relationship Between Spacing and Width Perception

RM trials produced both wider perceived widths and larger perceived spacing in the same participants (Figure 4B). The correlation between Δwidth and Δspacing was not significant (Figure 4C; r = 0.33, p = .158), suggesting that the RM-related increase in width perception was not reliably associated with the RM-related increase in perceived spacing across participants.

#### Array Extent

Perceived array extent was underestimated under redundancy masking. At every set size, the perceived extent was smaller (more underestimated) in RM than in no-RM trials (set size 3: β = 0.55, z = 12.81, p < .001, d = 0.84, 95% CI [0.47, 0.64]; set size 4: β = 0.55, z = 13.47, p < .001, d = 0.83, 95% CI [0.47, 0.63]; set size 5: β = 0.43, z = 12.01, p < .001, d = 0.66, 95% CI [0.36, 0.50]). Relative to the presented array, extent was underestimated in both conditions, but far more strongly under RM (no-RM: M = −0.18°, 95% CI [−0.29, −0.07], p = .001; RM: M = −0.69°, 95% CI [−0.81, −0.58], p < .001). Thus, even though perceived spacing was larger in RM trials, the net perceived extent of the array was reduced when items were lost, consistent with a radial underestimation of visual space (Figure 4D).

#### Density

Perceived density was largely unaffected by redundancy masking. The overall RM vs no-RM difference was small and not significant (β = −0.008, z = −1.26, p = .206, d = 0.06). Conditioned on set size, density differed significantly only at set size 3, where RM trials were perceived as slightly denser than no-RM trials (β = −0.028, z = −2.93, p = .003, d = 0.22, 95% CI [−0.047, −0.009]); no difference emerged at set size 4 (p = .85) or set size 5 (p = .52). Thus, despite the strong RM vs no-RM differences in the perceived number of bars, perceived width, and perceived spacing, the perceived density of the array remained very similar on RM and no-RM trials, changing far less than any of these measures (Figure 4E).

## DISCUSSION

The visual system receives far more information than it can process. Compression is one way of reducing this information. A central question is how compression changes the representation of the input: What information is lost, retained, or transformed? One way of compressing visual input is based on the extraction of redundancies, followed by a more compact representation. Redundancy masking is a prominent form of this compression: items in repeating patterns do not reach conscious awareness despite sufficient size and spacing between items well above the resolution limit (Sayim & Taylor, 2019; Yildirim et al., 2020, 2021, 2022). By presenting arrays of identical bars in the periphery, we found that participants frequently reported one bar fewer than presented, replicating redundancy masking. Importantly, the perceived number of bars was related to a systematic change in perceived width. On trials in which a bar was redundancy-masked (e.g., reporting two bars when three were presented), the perceived width of the bars was closer to the presented width than on trials in which all bars were reported. Hence, these results suggest that redundancy masking goes hand in hand with improved feature discrimination. Computational modeling separated the perceived width into the general peripheral underestimation (of width/size) and a contribution from the redundancy-masked bar, allowing us to estimate how much of the masked bar’s width was pooled into the perceived (remaining) bars. Most of the masked bar’s width was lost and only a small fraction contributed to the perceived width of the remaining bars. These findings suggest that redundancy masking reduces redundant information while representing the remaining information more accurately.

Redundancy is information in a signal that can be predicted from the rest of the signal, and its reduction is a fundamental principle of sensory coding (Alvarez & Oliva, 2009; Attneave, 1954; Barlow, 1961). Textures provide the canonical example: individual elements share information with their neighbors, allowing the visual system to represent the statistical regularities of a region without encoding every element individually (Julesz, 1981; Portilla & Simoncelli, 2000; Rosenholtz, 2014). Unlike textures, in which statistical regularities are defined across many elements, our displays instantiate the same principle of redundancy with only three to five elements. This makes it possible to examine the consequences of statistical representation at the level of individual items, including the loss of a single item from the percept. The bars in each array were identical, so each bar was fully predictable from its neighbors and carried no unique feature information. Redundancy masking could therefore reduce the number of repeated bars without eliminating access to their shared features: participants reported the shared feature of the array (width) highly accurately. This finding contrasts with the well-established detrimental effects of contextual interaction in peripheral vision (Bouma, 1970; Whitney & Levi, 2011; Herzog et al., 2015). When a target is flanked by nearby items, performance usually deteriorates—a phenomenon known as visual crowding, a major bottleneck of peripheral object recognition (Levi, 2008; Whitney & Levi, 2011). Contextual interactions, as in visual crowding, consistently impair the perception of individual targets. Relative improvements in target perception, by contrast, typically depend on specific conditions, such as long-grouping between remote identical items (Sayim et al., 2014; see also Yildirm-Keles et al., 2026) or the emergence of features that provide information about the target (Melnik et al., 2018; 2020). Redundancy masking is fundamentally different: all items are identical and the improvement requires no additional grouping or emergent structure. Instead, the improvement is directly coupled with the loss of an item from perception. Since RM and no-RM trials were physically identical displays, this coupling between losing an item and gaining feature accuracy must reflect a change in perception rather than a difference in stimulus. Importantly, this improvement also came at no cost to precision, without any reduction of the trial-to-trial consistency of the width judgments.

This improvement was evident in perceived width. On no-RM trials, the bars were perceived as narrower than their physical width in both experiments, consistent with the established underestimation of perceived size in peripheral vision (Newsome, 1972; Baldwin et al., 2016). The single-bar condition provided a baseline for peripheral width perception without any contextual interactions. Bars under RM resulted in width estimates that were at least as accurate as, or more accurate, than a bar presented alone: they matched the single bar in Experiment 2 and were closer to the presented width than the single bar in Experiment 1. Width perception on RM trials was therefore not merely at the level of a single bar, but more accurate than for a bar shown alone. This advantage over the single bar was not replicated in Experiment 2, where the single-bar baseline was already close to veridical, similar to performance on RM trials. Moreover, the main and baseline tasks in Experiment 2 were less directly comparable because the main task required participants to adjust both width and spacing rather than width alone. The main task therefore imposed more demands than the baseline task; nevertheless, width perception on RM trials still matched the baseline.

This pattern of results places an important constraint on current accounts of peripheral vision. Summary-statistic and averaging accounts propose that local features are pooled into a shared representation, such as the mean of the neighboring features (Parkes et al., 2001; Greenwood et al., 2009). However, averaging identical widths cannot alter their mean: whether one bar or several contribute to the estimate, the resulting width remains the same. Moreover, these accounts do not predict the loss of entire items when all items are identical. Population-code accounts provide a potential mechanism for item loss, because overlapping representations of similar items can become indistinguishable and yield fewer perceived items (e.g., van den Berg et al., 2010). However, this mechanism also does not predict a change in width depending on the perceived number of items. If identical bars are represented in the same neural population, their combined response takes an intermediate value, which, for identical bars, is simply their shared width. Hence the perceived number can decrease while the perceived width of the bars should not change. Here, we observed a coupling between item loss and feature perception that existing accounts do not predict: the feature representation of the surviving items changes systematically when another item is lost. In these accounts, perceived width does not depend on whether or not an item is redundancy-masked, and none predicts the wider—and more accurate— widths we observed.

We account for this coupling with a two-stage model: a peripheral width underestimation stage followed by a pooling stage. The first stage captures the general underestimation of peripheral width: each bar is represented with a multiplicative scaling factor below one, independent of redundancy masking. The second stage acts only when a bar is redundancy-masked, allowing a fraction of the width of an unreported bar to be pooled with the perceived bars. The pooling parameter (γ) sets how much of a redundancy-masked bar’s width is pooled with the perceived bars: when γ = 0 no pooling occurs and the bars retain the initial underestimated representation; when γ = 1, the entire width of the redundancy-masked bar is pooled, conserving its total width despite the reduction in the perceived number of bars. Intermediate values represent partial pooling. The data are consistent with weak pooling. For an array of three bars perceived as two, complete pooling would increase each perceived bar by 50%. Even after peripheral underestimation of the first stage, this would yield substantial overestimation of width. This was not observed: the perceived width in RM trials was close to veridical, with weak estimated pooling (γ = 0.20 in Experiment 1; γ = 0.32 in Experiment 2), only slightly counteracting peripheral underestimation of the first stage. Our model therefore suggests that most of the width of a redundancy-masked bar is lost with the item, while only a small fraction may be pooled with the perceived bars. In the model, the improved accuracy of RM trials is captured by the combination of general peripheral underestimation and partial compensation following redundancy masking.

Another possibility is that the improvement in width perception under redundancy masking is related to the larger perceived spacing on RM trials. In peripheral vision, contextual interactions between neighbouring items decrease with increasing spacing between them (Bouma, 1970; Toet & Levi, 1992; Pelli et al., 2004), with their strength determined by retinal as well as perceived separation (Herzog et al., 2015; Maus et al., 2011; Dakin et al., 2011). When perceived and retinal positions are dissociated, the strength of crowding follows the perceived positions (Maus et al., 2011; Dakin et al., 2011). Similarly, when items are physically present but do not reach awareness, performance is better predicted by the number of perceived items than by the number of presented items (Wallis & Bex, 2011). Together, these findings suggest that perceived spatial arrangement of a display can influence contextual interactions. In our displays, stimuli were physically identical in RM and no-RM trials, but under RM trials the perceived spacing between the remaining bars was larger. This larger perceived spacing may therefore have weakened contextual interactions and contributed to the more accurate width perception on RM trials.

Under redundancy masking, the perceived spacing between the remaining bars increased, while the overall array extent decreased. The reduction in array extent reflects the combined changes in the percept, including the loss of an item and the increase in spacing, rather than a change that can be attributed to any of these properties alone. Despite these systematic changes in perceived number, width, spacing, and array extent, array density (the proportion of the stimulus occupied by bars) was comparatively stable across RM and no RM trials. This preservation of density is compatible with summary-statistic accounts of peripheral vision. On these accounts, peripheral vision does not encode a list of individual items and their positions, but a set of summary statistics computed over regions that increase in size with eccentricity (Portilla & Simoncelli, 2000; Balas et al., 2009; Freeman & Simoncelli, 2011; Rosenholtz et al., 2019). A representation of this kind could preserve density even as the perceived item and position level properties of the array change, because density is a property of the whole array rather than of any individual item. However, existing summary-statistic models do not predict that items are lost from perception when they are identical. For example, the Texture Tiling Model (TTM; Balas et al., 2009; Keshvari & Rosenholtz, 2016) predicts uncertainty in the perceived number of items (Rosenholtz et al., 2019), but for identical, repeating arrays like ours, the TTM does not predict that an item is lost from perception (Keshvari & Rosenholtz, 2016, Fig. 5A). These models also do not predict the systematic local changes in appearance that accompanied the item loss in redundancy masking. Summary-statistics models represent images through summary statistics (Portilla & Simoncelli, 2000; Balas et al., 2009; Freeman & Simoncelli, 2011). Thus, a given set of summary statistics can be consistent with different local appearances. If the perceptual distortions observed were due solely to such a summary statistical representation, the local properties would not be expected to shift systematically in one direction. Instead, they could vary across the different local appearances compatible with the same pooled statistics. In our displays, however, the changes were consistently directional: perceived width and spacing increased, whereas array extent decreased under redundancy masking. This indicates that changes in local appearance associated with redundancy masking are systematic rather than variations distributed around the values determined by the stimulus,, and that appearance-based measures, like the adjustment task used here, are well suited to reveal this structure.

Compression allows the visual system to cope with an input that exceeds its limited capacity, and this compression is usually assumed to be lossy, reducing the amount of information at the expense of visual detail (Balas et al., 2009; Bates & Jacobs, 2020; Rosenholtz et al., 2012; Zhaoping, 2006). Our results show that the visual system can also exploit redundancy of the input during compression. When one item of a repeating array was redundancy-masked and did not reach awareness, the features of the perceived items were represented more accurately than when all items were perceived. Hence, the loss of items in peripheral vision can be accompanied by improved representation of the information that remains. More generally, redundancy masking suggests that compression in peripheral vision is not simply a loss of information but a mechanism that discards redundant information while preserving what is informative, with the features that remain represented more accurately.

## Data Availability

The raw and processed data generated in this study have been deposited in an Open Science Framework (OSF) repository and are available for review via an anonymized view-only link: https://osf.io/ek23j/?view_only=a870bc1def5a44588204b3073ad3b26d

## Code Availability

All analysis code (R) used in this study is available for review via an anonymized view-only link: https://osf.io/zva64/?view_only=8461f6380d894234a933b016a84add1f

## Acknowledgments

This study was supported by the Agence Nationale de la Recherche (ANR) under the Grant Number ANR-24-FRAL-0016-01, and the Scientific and Technological Research Council of Turkey (TUBITAK) under the Grant Number 122N748. Additional support by ANR-17-EURE-0017 Frontcog, ANR10-IDEX-0001-02 PSL.

