## Supplementary Material for "Compression of redundant visual information improves feature discrimination in human vision"

#### S1 Precision of width perception

For each participant, we quantified the precision of width perception as the standard deviation (SD) of width deviations, computed separately for each condition (RM trials, no-RM trials, and the single-bar control) and each presented width (0.25°, 0.4°, 0.55°). Because the variability of width judgments scales with the presented width, each width-specific SD was divided by its corresponding presented width, and the resulting values were averaged across widths, yielding one normalized precision value per participant and condition. Width cells with fewer than two trials, for which no SD can be computed, were omitted from this average; this affected one participant in Experiment 1, who had no RM trials at the 0.55° width, and whose RM precision was therefore averaged over the two remaining widths. Participants without a precision value in one or more of the three conditions were excluded from the paired comparisons, so that all three comparisons were based on the same set of participants. In Experiment 1, all 21 participants were retained. In Experiment 2, three participants were excluded: two lacked single-bar data entirely due to a coding error (IDs 701 and 703; see Section S3), and one (ID 702) completed only three single-bar trials (one per width), too few to compute an SD, leaving $N = 17$. Precision was compared across conditions using three paired $t$-tests (RM vs. no-RM, RM vs. single-bar, no-RM vs. single-bar), with $p$-values FDR-corrected.

The main-text precision measure was averaged across the three presented widths. Splitting it by presented width and analysing the RM and no-RM values with a linear mixed-effects model (precision $\sim$ RM type $\times$ bar width, with by-participant random effects) reproduced the width-averaged result. Precision improved for wider bars (main effect of width, both $p < .001$), but redundancy masking did not change it: neither the main effect of RM type nor the RM type $\times$ width interaction was significant (all $p \geq .09$), and no RM vs. no-RM contrast reached significance at any width in either experiment (all $p_{\text{FDR}} \geq .21$; Figure S1). The more accurate width perception on RM trials was thus not bought at the cost of noisier width estimates for any individual bar width.

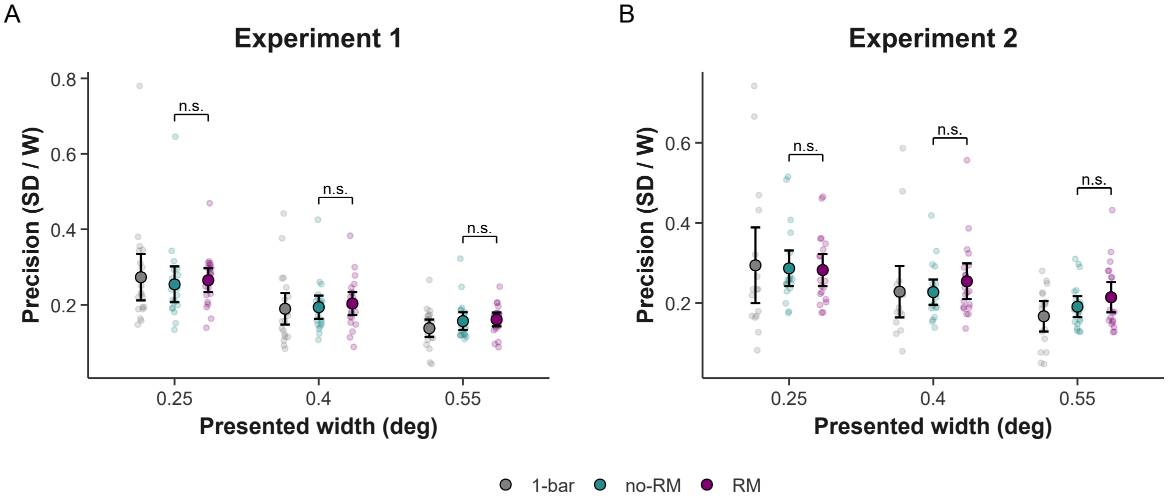


**Figure S1:** Precision of width perception by presented width. Precision is the standard deviation of raw width deviations divided by the presented width (smaller = more precise), split by presented width rather than averaged across widths as in the main text. (A) Experiment 1; (B) Experiment 2. Small points are individual participants; large points are group means with 95% confidence intervals for the single-bar baseline (grey), no-RM (teal), and RM (magenta) conditions. Brackets show RM versus no-RM `emmeans` contrasts from the linear mixed-effects model (FDR-corrected across the three widths), which were non-significant (n.s.) at every width in both experiments.

#### S2 Computational model specifications

To examine how perceived width depended on the number of perceived bars, we developed a family of computational models (M0–M5), each corresponding to a distinct hypothesis about the relationship between presented width and the perceived number of bars.

##### Null model (M0)

The starting point was the Null model (M0), which corresponds to the prediction of conventional averaging accounts: because all bars have the same width, averaging any subset of bars returns the same width, and perceived width should therefore remain unchanged regardless of how many bars are perceived (Parkes et al., 2001; Greenwood et al., 2009). M0 thus assumed that perceived width equals the presented width,

$$W' = W. \tag{1}$$

This model predicts neither underestimation nor any pooling of width information when bars are lost from observers' reports.

The remaining models were motivated by two established properties of peripheral vision. The first was peripheral size underestimation: overall perceived width is smaller than the presented width (Newsome, 1972), captured by a width-scaling parameter ($\alpha$). The second is pooling, which we extend beyond the conventional accounts by making pooling depend on the number of bars presented relative to the number perceived. For example, when fewer bars are perceived than presented, our model predicts that the width of the redundancy-masked bar is pooled into that of the remaining bars, captured by a pooling parameter ($\gamma$).

#### Perfect Pooling model (M1)

The Perfect Pooling model (M1) assumes that the width information of redundancy-masked bars is fully pooled into the remaining perceived bars, such that the sum of the total width information is preserved. That is, the sum of presented widths equals the sum of perceived widths,

$$N \cdot W = N' \cdot W', \tag{2}$$

where $N$ and $W$ are the number and bar width of the presented bars, and $N'$ and $W'$ are the number and bar width of the perceived (reported) bars. Solving for the perceived width gives

$$W' = \frac{N}{N'} \cdot W. \tag{3}$$

The ratio $N/N'$, the pool ratio, captures how many presented bars contribute to each perceived bar. On no-RM trials, the reported bar number is the same as the presented bar number ($N' = N$), so the pool ratio is 1 and perceived width equals presented width ($W' = W$). On RM trials, the reported bar number is less than the presented bar number ($N' < N$), the pool ratio exceeds 1, and perceived width ($W'$) is correspondingly larger. Reporting five bars as four, for example, gives a pool ratio of 5/4. Perceived width thus scales linearly with the pool ratio, and fewer perceived bars (when RM occurs) should each appear proportionally wider.

#### Partial Pooling model (M2)

The Partial Pooling model (M2) extended M1 by raising the pool ratio to the power of the pooling parameter ($\gamma$), allowing the strength of pooling to vary:

$$W' = \left(\frac{N}{N'}\right)^{\gamma} \cdot W. \tag{4}$$

The pooling parameter ($\gamma$) determines the strength of pooling: while the pool ratio ($N/N'$) specifies how much width is available for pooling, the pooling parameter specifies how much of it is actually pooled. When $\gamma = 1$, the model is equivalent to Perfect Pooling (M1); when $\gamma = 0$, the pool ratio drops out, and the model reduces to the Null model (M0). Intermediate values ($0 < \gamma < 1$) correspond to partial pooling, in which only a fraction of the redundancy-masked bar's width is pooled. Importantly, because the pool ratio exceeds 1 only on RM trials, $\gamma$ shapes width perception only when RM occurs, but has no effect on no-RM trials where the pool ratio is 1 and M2 again reduces to M0. Thus, M2 cannot capture peripheral size underestimation: bar widths are underestimated even when no bar is lost, and this underestimation requires a separate width-scaling parameter, introduced in the following models.

#### Width-scaling + Pooling model (M3)

The Width-scaling + Pooling model (M3) adds a width-scaling parameter ($\alpha$) to account for peripheral size underestimation combined with perfect (not partial) pooling:

$$W' = \alpha \cdot \frac{N}{N'} \cdot W. \tag{5}$$

When $\alpha < 1$, the model predicts that perceived width is compressed relative to the presented width. In principle, $\alpha$ can also exceed 1, which would correspond to an expansion rather than an underestimation of perceived width; given the well-established peripheral size underestimation, however, such values are highly unlikely. On no-RM trials (pool ratio = 1), the model reduces to $W' = \alpha \cdot W$. M3 can therefore capture peripheral size underestimation even in no-RM trials. On RM trials, the same width-scaling is applied, and the result then scales linearly with the pool ratio, i.e., perfect pooling of the redundancy-masked bars (M1). Thus, M3 tests whether perceived width can be explained by width-scaling and perfect pooling, while holding pooling strength constant.

#### Width-scaling + Partial Pooling model (M4)

The Width-scaling + Partial Pooling model (M4) combines both mechanisms, with a width-scaling parameter ($\alpha$) and a pooling parameter ($\gamma$):

$$W' = \alpha \cdot \left(\frac{N}{N'}\right)^{\gamma} \cdot W. \tag{6}$$

M4 dissociates two components of width perception: the width-scaling parameter ($\alpha$) captures the general tendency to perceive peripheral bars as thinner than they are, while the pooling parameter ($\gamma$) captures how much of the redundancy-masked bar's width is pooled into the remaining perceived bars (Figure 1B, 1C). M4 also contains the earlier models as special cases: when $\alpha = 1$ and $\gamma = 0$, it is equivalent to M0; when $\alpha = 1$ and $\gamma = 1$, it is equivalent to M1; when $\alpha = 1$, it is equivalent to M2; and when $\gamma = 1$, it is equivalent to M3.

#### Linear Width-scaling + Partial Pooling model (M5)

The Linear Width-scaling + Partial Pooling model (M5) extends M4 by letting the width-scaling parameter ($\alpha$) vary with bar width, replacing the constant $\alpha$ with a linear function of the presented width:

$$W' = \alpha(W) \cdot \left(\frac{N}{N'}\right)^{\gamma} \cdot W, \qquad \text{where } \alpha(W) = \alpha_0 + \alpha_1 W. \tag{7}$$

Here, $\alpha(W)$ captures the width-dependent width-scaling component, whereas $(N/N')^{\gamma}$ captures pooling as a function of the pool ratio. The intercept $\alpha_0$ is the hypothetical width-scaling at zero width, and the slope $\alpha_1$ represents the rate at which width-scaling changes with the presented width. Values of $\alpha(W) < 1$ indicate peripheral size underestimation and $\alpha(W) = 1$ indicates no width-scaling. On no-RM trials (pool ratio = 1), the model predicts $W' = \alpha(W) \cdot W$. On RM trials (pool ratio $> 1$), perceived width depends on both parameters: the pooling parameter ($\gamma$) determines how strongly the pool ratio increases perceived width, and the slope $\alpha_1$ determines whether this prediction rises or falls with bar width. Thus, M5 tests whether peripheral size underestimation scales with bar width while the pooling parameter is estimated independently. When $\alpha_1 = 0$, width-scaling is constant across widths, and M5 is equivalent to M4 with $\alpha = \alpha_0$.

##### Model fitting and comparison

All models were fit separately for each participant. For each model, parameters were estimated by minimizing the sum of squared errors between the reported width and the model-predicted width across trials:

$$\text{SSE} = \sum_{i=1}^{n} \left( W'_{\text{rep},i} - \hat{W}'_{\text{pred},i} \right)^2 . \tag{8}$$

Model comparison was conducted using two complementary approaches. First, the Akaike Information Criterion (AIC) was computed for each participant and model and used as the primary criterion for selecting the best fit-versus-complexity trade-off. Second, likelihood ratio tests (LRTs) were used to assess whether adding width-scaling or partial pooling significantly improved fit. Finally, for the best-fitting model, we examined participant-level parameters. We tested each participant's width-scaling parameter ($\alpha$) estimates against 1, since below 1 indicates peripheral size underestimation. For the pooling parameter ($\gamma$), we characterized its distribution relative to its two theoretical bounds. Because $\gamma = 0$ corresponds to no pooling, where the redundancy-masked bar's width is entirely lost, and $\gamma = 1$ corresponds to perfect pooling, where its width is fully pooled into the remaining bars, we classified each participant by whether their $\gamma$ fell below or above the midpoint of 0.5. Values below 0.5 indicate that more of the lost bar's width was lost than pooled into the bars that remained, and values above 0.5 the reverse.

#### S3 Two-stimulus discrimination control (Experiment 2)

Participants completed a two-stimulus discrimination control task to verify that the bars presented as stimuli could be spatially resolved at the eccentricities and spacings used in the experiments. On each trial, either one or two bars were presented for 150 ms, randomly to the left or right of fixation, at the farthest eccentricities of the main experiment (11.35° for 4-bar arrays, 11.8° for 3- and 5-bar arrays). Participants indicated whether they perceived one or two bars. For 2-bar arrays, the widths (0.25°, 0.4°, 0.55°) and spacings (0.7°, 0.9°, 1.1°) were identical to those used in the main experiment. This control ensured that the stimuli remained above the participants' visual resolution limits. The task included 18 unique trials (2 set sizes $\times$ 3 widths $\times$ 3 spacings) repeated 6 times, yielding 108 trials in total.

In Experiment 2, a coding error left two participants (IDs 701 and 703) without single-bar baseline data, so single-bar analyses in Experiment 2 use the remaining 18 participants. These same two participants also lacked two-stimulus discrimination data, which was therefore available for 18 of the 20 participants. Accuracy was well above the 50% chance level for every participant (Figure S2), with a mean of 95.4% ($SD = 4.9$) and a range of 80.6% to 100%. These results confirm that the bars were spatially resolvable at the eccentricities and spacings used in the main experiment.

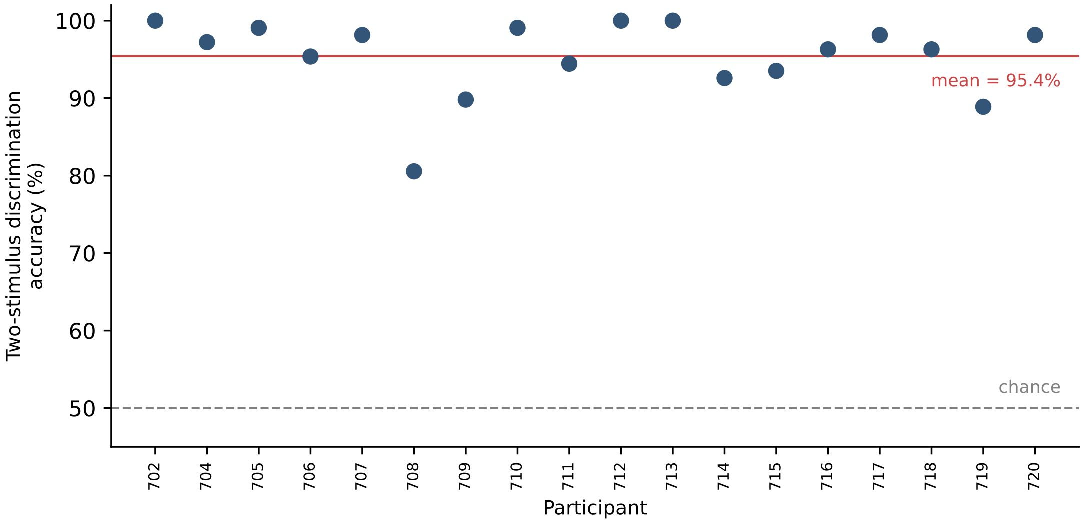


**Figure S2:** Two-stimulus discrimination accuracy in the Experiment 2 control task. Each point is one participant's percentage of correct one-versus-two-bar judgments. The dashed line marks chance (50%) and the red line the group mean (95.4%). Participants 701 and 703 lacked control data (see text).

### S4 Supplementary Tables

**Table S1:** Distribution of number deviations, Experiments 1 and 2. Number deviation is the reported minus the presented number of bars. Counts are pooled across participants; percentages are relative to all trials retained after the response-time and ±4 number deviation exclusions (see main text, Data analysis). Analyses comparing RM and no-RM trials were restricted to number deviations of $-1$ (RM) and 0 (no-RM); multi-bar losses (deviations $\leq -2$) and over-reports (deviations $\geq +1$) were excluded.

| | Exp. 1 ($N = 21$) | | Exp. 2 ($N = 20$) | |
|---|---|---|---|---|
| Number deviation | Trials | % | Trials | % |
| $-4$ | – | – | 1 | 0.02 |
| $-3$ | 8 | 0.07 | 2 | 0.03 |
| $-2$ | 352 | 3.23 | 101 | 1.63 |
| $-1$ (RM) | 3460 | 31.71 | 2057 | 33.15 |
| 0 (no-RM) | 6143 | 56.30 | 3651 | 58.83 |
| +1 | 776 | 7.11 | 353 | 5.69 |
| +2 | 156 | 1.43 | 40 | 0.64 |
| +3 | 16 | 0.15 | – | – |
| +4 | – | – | 1 | 0.02 |
| Multi-bar losses ($\leq -2$) | 360 | 3.30 | 104 | 1.68 |
| Over-reports ($\geq +1$) | 948 | 8.69 | 394 | 6.35 |
| Total excluded | 1308 | 11.99 | 498 | 8.02 |
| Total trials | 10911 | 100 | 6206 | 100 |

**Table S2:** Percentage of redundancy-masking (RM) trials by condition, Experiments 1 and 2. Values are participant means (*SD*). RM% is the percentage of RM trials (number deviation = −1) out of RM and no-RM trials (number deviation = −1 or 0), computed per participant and averaged. Spacing categories (Experiment 2 only) were formed by splitting the range of presented inter-bar spacings into tertiles.

| | | RM% *M* (*SD*) | |
|---|---|---|---|
| Set size | Bar width (°) | Exp. 1 ($N = 21$) | Exp. 2 ($N = 20$) |
| 3 | 0.25 | 28.4 (25.6) | 21.2 (20.1) |
| 3 | 0.40 | 27.8 (22.3) | 19.7 (22.0) |
| 3 | 0.55 | 29.2 (26.5) | 21.3 (22.0) |
| 4 | 0.25 | 34.0 (24.5) | 27.0 (20.8) |
| 4 | 0.40 | 37.7 (22.7) | 30.6 (22.0) |
| 4 | 0.55 | 46.1 (25.5) | 37.7 (21.2) |
| 5 | 0.25 | 40.0 (27.9) | 51.6 (29.4) |
| 5 | 0.40 | 40.9 (29.0) | 53.9 (29.5) |
| 5 | 0.55 | 45.5 (27.7) | 58.5 (26.8) |
| Set size | Spacing (Exp. 2) | | |
| 3 | Smaller | – | 24.2 (24.0) |
| 3 | Middle | – | 25.1 (24.5) |
| 3 | Larger | – | 18.9 (21.0) |
| 4 | Smaller | – | 36.1 (21.2) |
| 4 | Middle | – | 28.3 (23.1) |
| 4 | Larger | – | 24.3 (23.6) |
| 5 | Smaller | – | 60.5 (29.1) |
| 5 | Middle | – | 49.6 (28.7) |
| 5 | Larger | – | 48.2 (28.4) |

**Table S3:** Model comparison, Experiments 1 and 2. $k$ = number of free parameters; mean AIC and mean RMSE are computed per participant and averaged (lower indicates better fit); best-fit ($n$) is the number of participants for whom the model had the lowest AIC.

| Model | $k$ | Mean AIC | Mean RMSE | Best-fit ($n$) |
|---|---|---|---|---|
| *Experiment 1* ($N = 21$) | | | | |
| M0: Null | 0 | −841.93 | 0.099 | 0 |
| M1: Perfect Pooling | 0 | −549.11 | 0.142 | 0 |
| M2: Partial Pooling | 1 | −891.34 | 0.093 | 1 |
| M3: Width-scaling + Pooling | 1 | −914.86 | 0.092 | 0 |
| M4: Width-scaling + Partial Pooling | 2 | −1025.97 | 0.082 | 2 |
| M5: Linear Width-scaling + Partial Pooling | 3 | −1085.59 | 0.077 | 18 |
| *Experiment 2* ($N = 20$) | | | | |
| M0: Null | 0 | −448.69 | 0.113 | 1 |
| M1: Perfect Pooling | 0 | −312.37 | 0.142 | 0 |
| M2: Partial Pooling | 1 | −468.26 | 0.108 | 1 |
| M3: Width-scaling + Pooling | 1 | −487.84 | 0.105 | 0 |
| M4: Width-scaling + Partial Pooling | 2 | −531.17 | 0.097 | 3 |
| M5: Linear Width-scaling + Partial Pooling | 3 | −559.18 | 0.092 | 15 |

**Table S4:** Likelihood ratio tests for nested model comparisons, Experiments 1 and 2. Group $\chi^2$ is the sum of per-participant likelihood-ratio statistics ($df = 1$ each; group $df$ equals the number of participants). Sig. ($n/N$) is the number of participants for whom the added parameter significantly improved fit ($p < .05$).

| Comparison | Added parameter | $\chi^2$ | $df$ | $p$ | Sig. ($n/N$) |
|---|---|---|---|---|---|
| *Experiment 1* | ($N = 21$) | | | | |
| M0 vs. M2 | $\gamma$ (pooling) | 1079.59 | 21 | $< .001$ | 18/21 |
| M1 vs. M3 | $\alpha$ (width-scaling) | 7722.86 | 21 | $< .001$ | 21/21 |
| M2 vs. M4 | $\alpha$ added to $\gamma$ | 2869.25 | 21 | $< .001$ | 18/21 |
| M3 vs. M4 | partial vs. perfect $\gamma$ | 2375.33 | 21 | $< .001$ | 21/21 |
| M4 vs. M5 | width-dependent $\alpha$ | 1293.89 | 21 | $< .001$ | 16/21 |
| *Experiment 2* | ($N = 20$) | | | | |
| M0 vs. M2 | $\gamma$ (pooling) | 431.44 | 20 | $< .001$ | 14/20 |
| M1 vs. M3 | $\alpha$ (width-scaling) | 3549.27 | 20 | $< .001$ | 19/20 |
| M2 vs. M4 | $\alpha$ added to $\gamma$ | 1298.20 | 20 | $< .001$ | 16/20 |
| M3 vs. M4 | partial vs. perfect $\gamma$ | 906.58 | 20 | $< .001$ | 19/20 |
| M4 vs. M5 | width-dependent $\alpha$ | 600.25 | 20 | $< .001$ | 15/20 |

**Table S5:** Parameter estimates for the best-fitting model (M5), Experiments 1 and 2. Values are group means ($SD$). $\alpha_0$ and $\alpha_1$ are the intercept and width-dependent slope of the width-scaling function $\alpha(W) = \alpha_0 + \alpha_1 W$; the effective width-scaling $\alpha(W)$ is given for each bar width; $\gamma$ is the pooling parameter.

| | | | Effective $\alpha(W)$ | | | | |
|---|---|---|---|---|---|---|---|
| Experiment | $\alpha_0$ | $\alpha_1$ | 0.25° | 0.40° | 0.55° | $\gamma$ | $\gamma < 0.5$ |
| Experiment 1 ($n = 21$) | 1.03 (0.27) | −0.32 (0.55) | 0.95 | 0.90 | 0.86 | 0.20 (0.12) | 21/21 |
| Experiment 2 ($n = 20$) | 1.02 (0.28) | −0.30 (0.67) | 0.95 | 0.90 | 0.86 | 0.32 (0.22) | 17/20 |
